# The Nβ motif of NaTrxh directs secretion as an endoplasmic reticulum transit peptide and variations might result in different cellular targeting

**DOI:** 10.1101/2023.05.31.543037

**Authors:** Andre Zaragoza-Gómez, Emilio García-Caffarel, Yuridia Cruz-Zamora, James González, Víctor H. Anaya-Muñoz, Felipe Cruz-García, Javier Andrés Juárez-Díaz

## Abstract

Soluble secretory proteins with a signal peptide reach the extracellular space through the endoplasmic reticulum-Golgi conventional pathway. During translation, the signal peptide is recognised by the secretory recognition particle and results in a co-translational translocation to the endoplasmic reticulum to continue the secretory pathway. However, soluble secretory proteins lacking a signal peptide are also abundant, and several unconventional (endoplasmic reticulum/Golgi independent) pathways have been proposed and some demonstrated. This work describes new features of the secretion signal called Nβ, originally identified in NaTrxh, a plant extracellular thioredoxin, that does not possess an orthodox signal peptide. We provide evidence that other proteins, including thioredoxins type *h*, with similar sequences are also signal peptide-lacking secretory proteins. To be a secretion signal, positions 5, 8 and 9 must contain neutral residues –a negative residue in position 9 in animal proteins– to maintain the Nβ motif negatively charged and a hydrophilic profile. Moreover, our results suggest that the NaTrxh translocation to the endoplasmic reticulum occurs as a post-translational event. Finally, the Nβ motif sequence at the N- or C-terminus could be a feature that may help to predict protein localisation, mainly in plant and animal proteins.

## Introduction

Protein secretion is an important cellular mechanism essential for many processes in all organisms, such as cell division and proper response to biotic and abiotic stresses [1,2]. In plant cells, for instance, the cell wall contains components, such as glycoproteins and enzymes, some of which are integrated into the plasma membrane (PM) and others are secreted towards the extracellular space to form part of the cell wall itself [3–5]. These components predominantly reach the PM via the vesicular trafficking endomembrane system.

It is thought that both membrane-integral components and secreted ones are localised via the well-characterised conventional protein secretion route. This pathway involves proteins that contain a signal peptide (SP), mainly characterised by its N-terminal position and its hydrophobic profile [1,6,7]. This is recognised by the signal recognition particle (SRP) –a conserved ribonucleoprotein– [1,8]. When an SP-containing protein is being translated, the SRP recognizes the SP and transports the mRNA-ribosome-nascent protein complex towards the endoplasmic reticulum (ER), where the SP is cleaved, and translation fully occurs [8]. Then, the protein is transported within vesicles to the Golgi apparatus, from which vesicles are released towards the PM. Once the vesicles fuse to the PM, the secretory proteins are released to the extracellular space whilst membrane-integral components remain as part of new PM material [1,5,9].

The conventional secretory pathway allows for important post-translational modifications required for specific protein functions and/or solubilisation once in the extracellular space. It is in the ER and mainly in the Golgi apparatus where protein glycosylation and oligosaccharide formation occur, which in plant cells is essential for cell wall deposition [3,5].

Reports regarding SP-lacking or leader-less secreted proteins have been increasing for a long time. This means the occurrence of secretory pathways that are ER/Golgi independent, referred to as the unconventional protein secretory routes (either ER, Golgi, or ER-Golgi independent are considered) [10,11]. All these ER/Golgi independent pathways – some hypothesised and others experimentally verified– direct soluble leader-less proteins towards the extracellular space and have been mainly studied in mammalian and yeast cells, although in plants, they have gained increasing relevance since the list of SP-lacking secreted plant proteins has been growing [2,10–12].

One such SP-lacking secretory protein is the thioredoxin (Trx) NaTrxh, identified in *Nicotiana alata*, that localises to the extracellular matrix of stylar transmitting tissue [13]; it is clustered within subgroup 2 of type *h* Trxs (Trxh-S2), which share an N-terminal extension –some also possess a C-terminal one– with sequences which are not fully conserved [13,14]. *In vitro* NaTrxh interacts with and reduces the S-RNase protein [13,15,16], which is a secretory protein of the stylar transmitting tissue and is the female S-determinant of the S-RNase-based self-incompatibility (SI) system (rev. within [17–19]). S-RNase reduction by NaTrxh results in a seven-fold increase in ribonuclease activity that is essential for self-pollen rejection in *Nicotiana* [16].

Different algorithms used to predict cellular localisation produce contradictory results regarding NaTrxh cellular localisation [13]. The first 16 amino acid residues of NaTrxh (Met-1 to Ala-16, referred to as the Nα motif; Fig 1A) do not participate in its secretion [15], as predicted by *hidden Markov* models [13,20]. Instead, the sequence between Ala-17 and Pro-27 –denominated as the Nβ motif (Fig 1A)– directs NaTrxh secretion, despite not being at the very N-terminus and possessing a predominantly hydrophilic profile [15]. This suggested that SRP should not be able to recognise the Nβ motif and therefore, NaTrxh secretion would follow an unconventional secretion pathway. However, when immunolocalised in *N. alata* styles, NaTrxh was associated with membrane systems, including the ER, the Golgi apparatus and presumably secretion vesicles [15].

**Fig 1.**
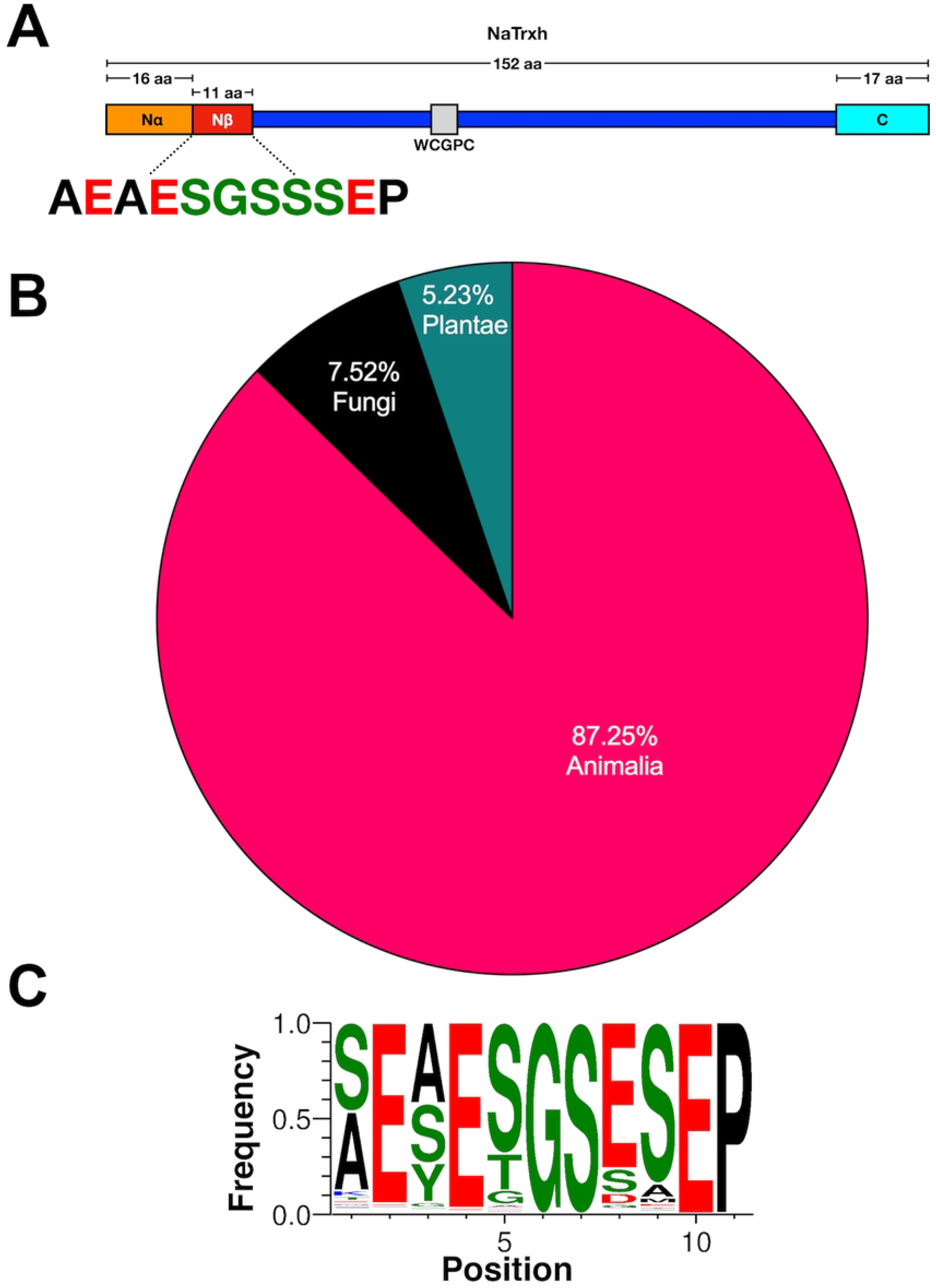
The Nβ motif sequence is found in proteins from all taxa. (A) Sequence of the Nβ motif in NaTrxh. The colour code refers to the charge of each amino acid (green: neutral; black: hydrophobic; red: negative). (B) Distribution of the eukaryotic proteins that contain the Nβ motif or a similar one. (C) Consensus sequence (featured as logos) of all the Nβ motifs found in eukaryotic proteins. Colour code as in (A).

The fact that NaTrxh secretion is due to the Nβ motif was proven by fusing its short sequence to the N-terminal of the green fluorescent protein (GFP) –creating an Nβ-GFP fusion protein– and tested by transient expression assays in onion epidermal cells. NaTrxh-GFP and Nβ-GFP fusion proteins were secreted using the conventional secretion elements: both passed through the ER and secretion was brefeldin-A sensitive –*i.e.*, proceeded via the Golgi apparatus– [15].

While soluble secretory SP-containing proteins are considered to follow the conventional secretory pathway, those that do not contain an orthodox SP are suggested to be secreted by an ER/Golgi independent one. However, through an analysis of the Nβ motif from NaTrxh, we hypothesised that NaTrxh follows a “semi-conventional” secretion pathway in which the Nβ motif functions as a transit peptide towards the ER to continue its secretory pathway to reach the extracellular space. This suggests a post-translational translocation instead of a co-translational one, as occurs with a typical SP-containing protein recognised by the SRP when translated. In addition, our data suggest that the Nβ motif might help to predict whether a protein lacking an SP might be secreted, specifically in plant proteins and presumably in animal ones. Mainly, among Trxh-S2 with similar N-extensions, two major groups were found, one with a similar Nβ motif that could be predicted to act as secretory proteins and another that may be a mitochondrial one, suggesting that few changes within this the Nβ motif might direct proteins towards different cellular compartments, contributing to the vast physiological roles in which Trxs are being involved.

## Results

### The eleven-residue long motif, Nβ, is present in transport and cell traffic proteins

The Nβ motif sequence (Fig 1A), identified as essential for NaTrxh secretion in plant cells [15], was used for BLASTP analysis. The output raised 383 protein sequences containing identical or similar sequences, 304 of them being eukaryotic proteins (S1 Table), all registered at the National Center for Biotechnology Information database (NCBI; http://www.ncbi.nlm.nih.gov). Within the eukaryotic protein sequences containing an Nβ motif (or similar), 266 are from animals, 22 from fungi and 16 from plants (Fig 1B and S1 Table). All 304 sequences were used to generate the Nβ motif-like eukaryotic consensus sequence shown in Fig 1C.

The small size of the Nβ motif itself, just eleven amino acid residues long, may lead to random identification of similar but functionally-unrelated sequences that would not represent any relationship to a secretory signal as hypothesised. However, 191 of the animal proteins detected are associated with membrane traffic, from which 98 are involved in endocytosis and exocytosis; among the plant proteins found, four are identified as chaperones belonging to the *Nicotiana* genus and annotated as Trxh-S2, where NaTrxh is grouped (S1 Table) [13]. The chance that these results were due to random factors is diminished because most proteins containing the Nβ motif of NaTrxh –or a similar one– are related to membrane-traffic mobility in cellular processes, suggesting an Nβ presence-function relationship.

### Amino acid positions in the Nβ motif required for secretion in plant cells

The consensus sequence of the Nβ motif comprising all sequences found in the different eukaryotic proteins (Fig 1C) showed that positions 1 and 3 are the least conserved and include two hydrophobic amino acid residues (Figs 1A and 1C). To determine if these amino acids were functionally relevant and to define the minimum size of the functional Nβ motif, we generated two types of protein variants (Fig 2A). The first one used the NaTrxh-GFP fusion protein, to which deletions of different sizes at its N-terminus were made (single residues at either 3 or 6 positions from the Nα motif): NaTrxhΔNα(+3)-GFP and NaTrxhΔNα(+6)-GFP; the second type was generated by deleting 3 or 6 amino acid residues from start of the Nβ-GFP sequence: Nβ(-3)-GFP and Nβ(-6)-GFP. All these protein variants were transiently expressed in onion epidermal cells.

**Fig 2.**
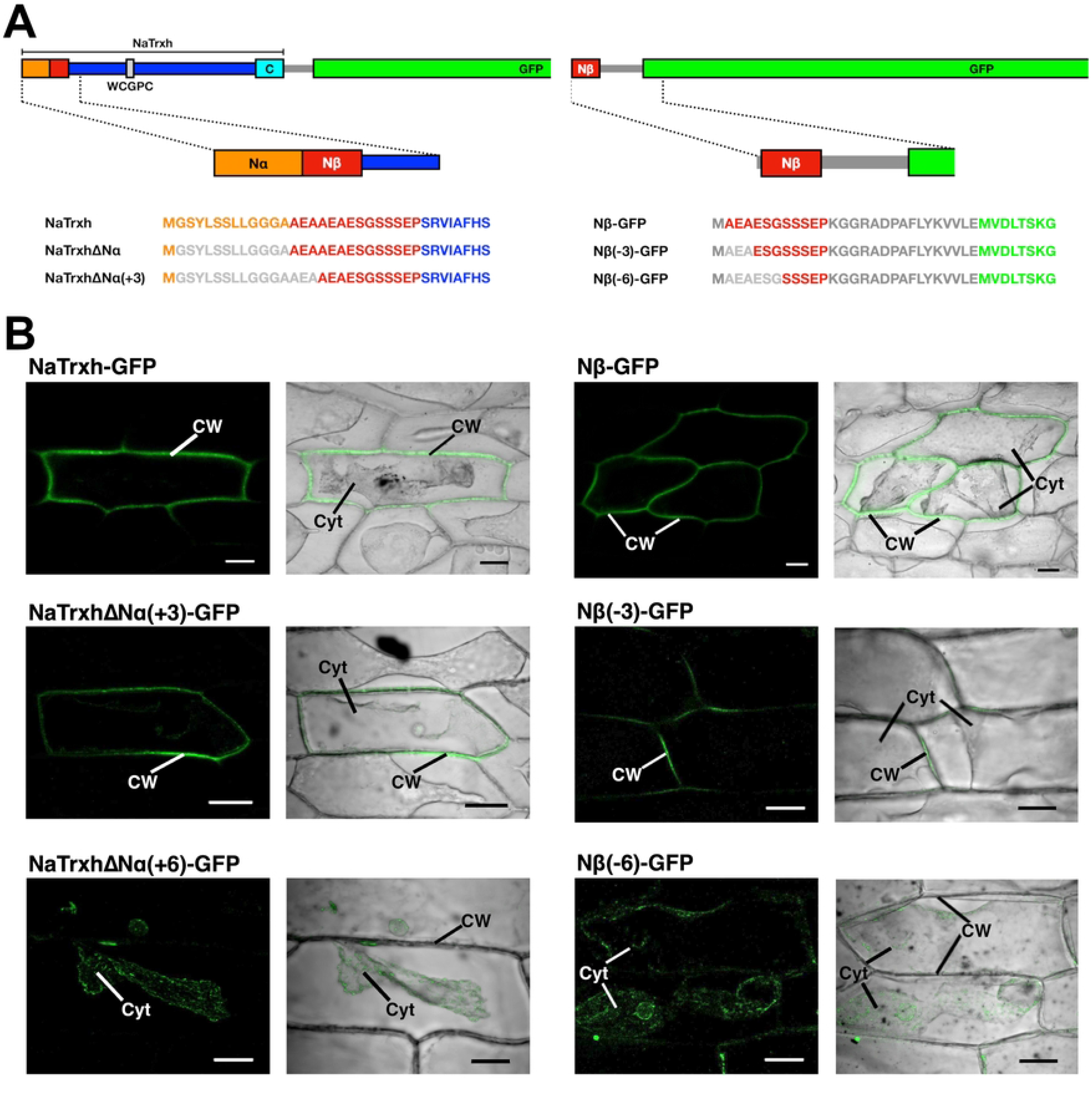
The first three Nβ motif residues do not contribute to its secretion signal function. (A) Schematic representation of the two types of constructs generated to assess the size of the functional Nβ motif. In one, deletions were made from NaTrxh fused to GFP (left) and the other from the Nβ motif directly fused to GFP (right). (B) Transient expression assays in onion epidermal cells and observed under confocal microscopy for GFP fluorescence to determine the localisation of NaTrxh-GFP, Nβ-GFP, NaTrxhΔNα(+3)-GFP, Nβ(-3)-GFP, NaTrxhΔNα(+6)-GFP, Nβ(-6)-GFP. Left panels: GFP fluorescence; right panels: merged image of bright field plus GFP fluorescence. Cells were pre-incubated with 1 M NaCl for plasmolysis. Cyt: cytoplasm; CW: cell wall. Scale bar: 50 μm.

When the Nβ deletions were transiently expressed in onion epidermal cells, we found that the first three positions are not required for the Nβ motif to direct protein secretion because while both NaTrxhΔNα(+3)-GFP and Nβ(-3)-GFP were localised in the extracellular space, NaTrxhΔNα(+6)-GFP and Nβ(-6)-GFP localised to the cytoplasm (Fig 2B). NaTrxh-GFP and Nβ-GFP, used as controls, both localised to the extracellular space (Fig 2B) as previously reported by [15]. These results confirm that the Nβ motif-guided secretion is independent of the core of NaTrxh, as previously reported [15].

As indicated above, the consensus sequence of the Nβ motif shown in Fig 1C is formed from all the eukaryotic sequences found in the set of predicted secretory and cytoplasmic proteins obtained from the BLASTP search (S1 Table). To discriminate between these groups, we used the *UniProtKB* database [21], where the information regarding cellular localisation is based on a score that varies from low information (score 1) to complete and experimental evidence (score 5). We clustered these proteins into four categories (from *A* to *D*) based on *UniProtKB* scores and their cellular localisation (Fig 3A): *A*, secreted proteins with score=5 (8 sequences); *B*, secreted proteins with score over 1 and under 3 (78 sequences); *C*, cytoplasmic proteins with score=5 (8 sequences); *D*, proteins with no information regarding their cellular localization (210 sequences).

**Fig 3.**
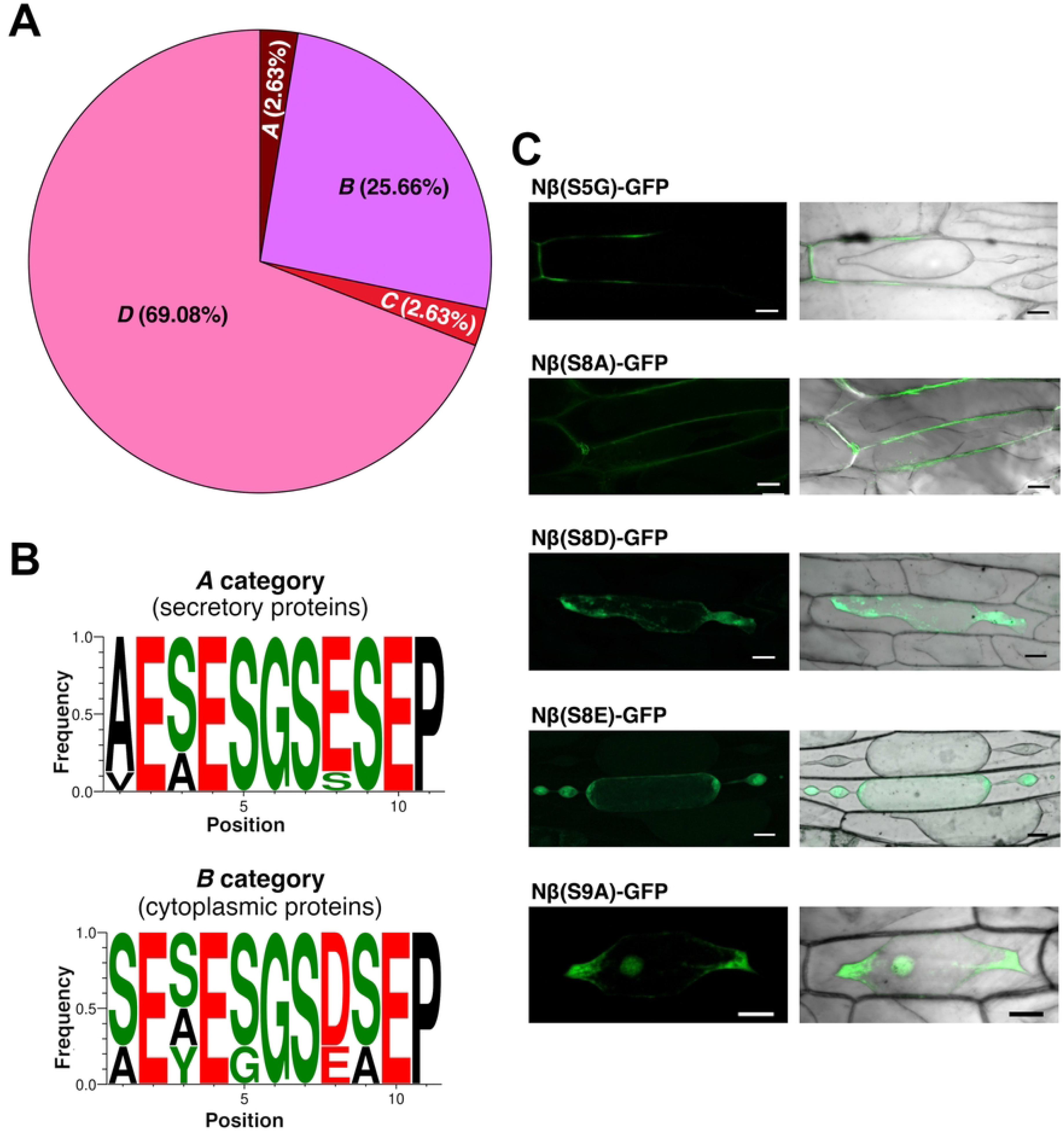
Positions 5, 8 and 9 within the Nβ motif differ between secretory and cytosolic proteins. (A) Distribution of the protein sequences among the categories generated based on the *UniProtKB* database regarding cellular localisation: *A* category (2.63%); *B* category (25.66%); *C* category (2.63%); *D* category (69.08%). (B) Logos of the sequences found in *A* category (upper) and *C* category proteins (down). (C) Transient expression assays in onion epidermal cells (as in Fig 2) of the different Nβ variants fused to GFP. Cyt: cytoplasm; CW: cell wall. Scale bar: 50 μm.

Although there is ample evidence regarding NaTrxh secretion [13,15], NaTrxh was grouped within the *D* category with other Trxh-S2 that have the same N-terminal extension sequence as NaTrxh. Based on the *UniProtKB* available data, this finding means the *A* category is likely underestimated. The *D* category appears to be overrated –it is indeed the largest category (Fig 3A)–. The same may occur with the *B* category, for which finding an Nβ motif might predict protein secretion, increasing its score (as with the *A* category) as discussed below. The results shown in Fig 3A and further analysis consider NaTrxh to reside within the *A* category.

To assess which amino acid residues might be essential for the Nβ motif to work as an SP-like sequence, we compared the two distinctive consensus sequences generated by contrasting localization-category proteins (featured as logos; Fig 3B): (1) generated from *A* category proteins; and (2) from *C* category proteins. Comparing these two consensus sequences revealed that positions 5, 8 and 9 vary from one group to another. While the *A* category proteins contain a serine residue in positions 5 and 9, some *C* category ones contain Gly-5 and Ala-9. Thus, to test whether these changes are relevant for the Nβ motif to direct secretion, we generated the Nβ(S5G) and the Nβ(S9A) variants –both replacing an *A* category residue with a *C* category residue– using the *Nβ-GFP* sequence as a template, which contains the NaTrxh Nβ sequence (Fig 1A). The results from transient expression in onion epidermal cells showed that while Nβ(S5G)-GFP was secreted, Nβ(S9A)-GFP was localised within the cytoplasm (Fig 3C). This provides evidence that, while variations in position 5 of the Nβ motif appear not to be relevant, position 9 requires a serine residue (or a neutral one) to maintain its role as a secretion signal. Therefore, the motif cannot be predicted to act as a secretion signal sequence if an Ala-9 –or probably any hydrophobic residue at this position– is present.

Regarding position 8, while most of the category *A* proteins contain a glutamate residue –NaTrxh being an exception since it contains a serine, which is considered in the consensus sequence of Fig 3B–, the *C* category proteins mainly contain an aspartate residue here and only a few have glutamate at this position (Fig 3B). Therefore, we generated the following variants fused to GFP: (1) Nβ(S8D) to mimic the sequence found in proteins within *C* category; (2) Nβ(S8E), which is the predominant form in the *A* category and could be expected to work as a signal peptide-like sequence; and (3) Nβ(S8A) to assess the relevance of a negatively charged residue at this position.

Results (Fig 3C) indicated that the protein variant Nβ(S8A)-GFP is localised at the extracellular space (Fig 3C), indicating that the change from a neutral to a hydrophobic residue did not affect the role of the Nβ motif as a secretion signal at this position. However, when Ser-8 was replaced by a negative residue as found in the Asp and Glu residues, both Nβ(S8D)-and Nβ(S8E)-GFP were localised within the cell (Fig 3C). These data provide different possible scenarios. First, Ser-8 appears essential for the Nβ motif to direct protein secretion. However, the amino acid predominantly found at this position is Glu in secretory proteins (*A* category). Notably, as Glu-8 was found in animal proteins, a glutamate at this position is suggested to be crucial for the Nβ motif to work as a secretion signal within this taxon. This would also explain why Asp-8 was found in cytoplasmic proteins (*C* category). In contrast, plant proteins appear to require a serine at position 8. This hypothesis is reinforced by the fact that NaTrxh was grouped within the *D* category (proteins with no information regarding its cellular localization) according to the available *UniProtKB* data despite the clear evidence supporting its secretion [13,15]. Some proteins within the *D* category are annotated as Trxh-S2, meaning that they have N-terminal extensions and, notably, they all possess the Ser-8 residue (Fig 4), reinforcing the hypothesis that the *D* category –and probably also *B*– might be overrated, and *A* category underrated.

**Fig 4.**
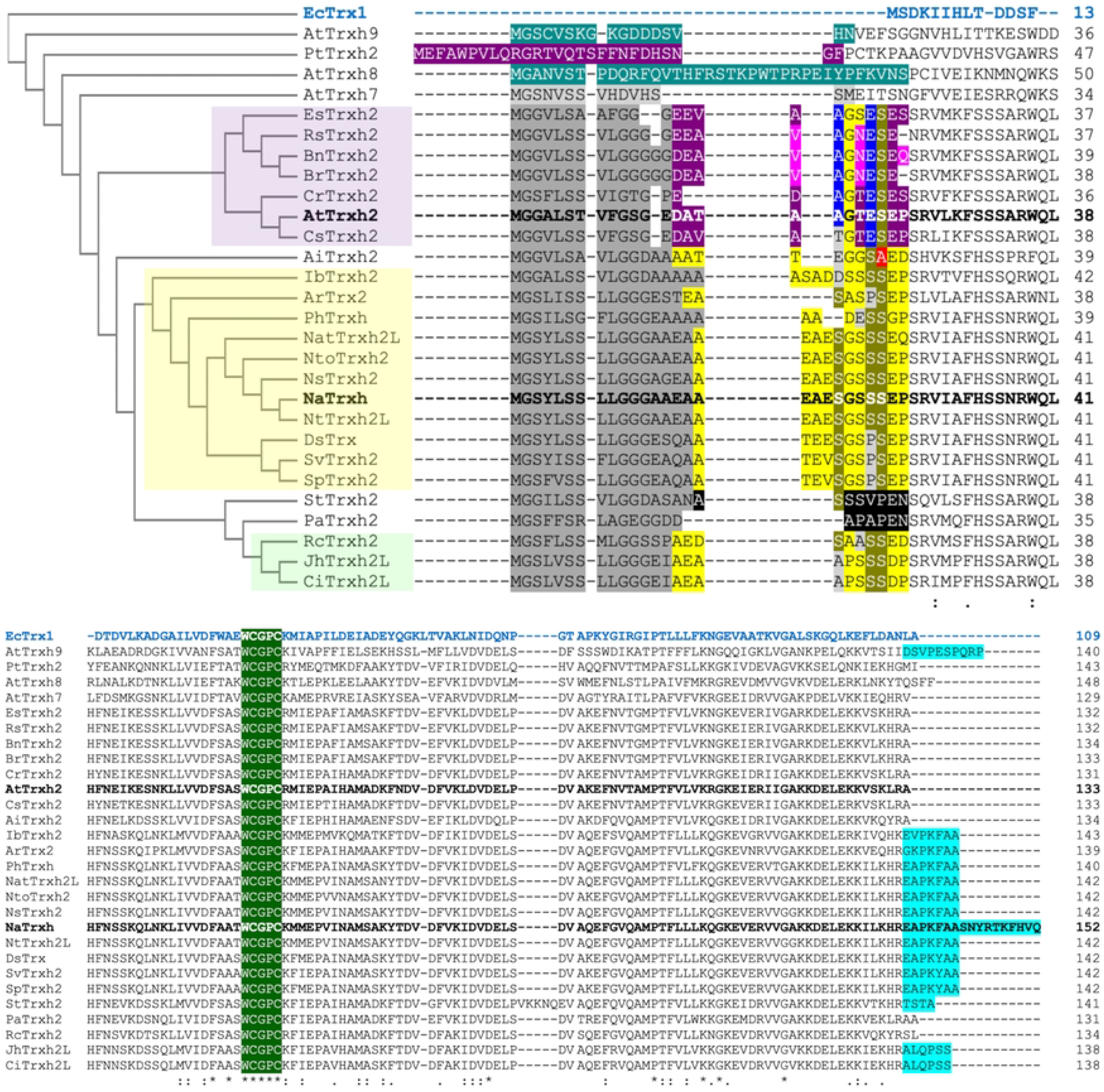
Analysis of different N-extensions of Trxh-S2. Multiple alignment analysis of different Trxh-S2 considering *E. coli* Trx1 (EcTrx1; AAA24693.1; blue sequence) as an outgroup. All Trxh-S2 are longer than *E. coli* Trx1, predominantly due to the N-terminal extensions present in all Trxh-S2 sequences and the C-terminal extensions present in some of them (cyan). Notably, NaTrxh (AAY42864.1; bolded) possesses the longest C-terminal extension. The extensions of those Trxs that have been associated to the plasma membrane are labelled in turquoise. Dark grey areas: Nα motifs or equivalent. Yellow: Nβ sequences; purple: Nβ variations that might result in a mitochondrial transit peptide; black: shortened Nβ sequences. The relevant identified differences are highlighted in blue. The sequences used and their respective GenBank accession codes are: *A. thaliana*: AtTrx2 (NP_198811.1); AtTrxh2 (Q38879.2; in bold for its mitochondrial localisation), AtTrxh7 (AAD39316.1), AtTrxh8 (AAG52561.1), AtTrxh9 (AAS49091.1); *Populus* hybrid: PtTrxh2 (AAL90749.1); *Eutrema salsugineum*: EsTrxh2 (XP_024014606.1); *Raphanus sativus*: RsTrxh2 (XP018446242.1); *Brassica napus*: BnTrxh2 (XP_013647760.2); *B. rapa*: BrTrxh2 (009125122.1); *Capsella rubella*: CrTrxh2 (XP006284753.1); *Camelina sativa*: CsTrxh2 (XP_019089899.1); *Arachis ipaensis*: AiTrxh2 (016194214.1); *Ipomea batatas*: IbTrxh2 (AAQ23133.1); *Actinidia rufa*: ArTrx2 (GFY87055.1); *Petunia hybrida*: PhTrxh (UYX45859.1); *Nicotiana attenuata*: NatTrxh2L (XP_019242743.1); *N. tomentosiformis*: NtoTrxh2 (XP_009602293.1); *N. sylvestris*: NsTrxh2 (XP_009795987.1); *N. tabacum*: NtTrxh2L (XP016508831.1); *Datura stramonium*: DsTrx (MCD9559198.1); *Solanum verrucosum*: SvTrxh2 (XP049346357.1); *S. pennellii*: SpTrxh2 (XP_015063846.1); *Senna tora*: StTrxh2 (KAF7819668.1); *Prosopis alba*: PaTrxh2 (XP_028806431.1); *Rosa chinensis*: RcTrxh2 (XP_024180631.1); *Juglans* hybrid: JhTrxh2L (XP_041003776.1); *Carya illinoinensis*: CiTrxh2L (XP_042969725.1).

### The Nβ motif might be crucial for targeting of Trxs to different cellular locations

In plants, at least eight types of Trxs have been identified, and there is a relationship between cellular localisation and Trx function (reviewed in [22]). For example, Trx types *f*, *m*, *x*, and *y* are chloroplast proteins, type *o* Trxs localise to the mitochondria, and those of type *s* are associated with the ER [23–28]. Type *h* Trxs (Trxh) form the largest and most heterogenous group and have been considered as cytosolic proteins [14,29]. However, there is evidence of some exceptions, like PtTrx*h*2 from *Populus tremula* and Trx*h*-2 from *A. thaliana* (AtTrxh2), which localise to the mitochondria [30–31] or NaTrxh, which is located at the extracellular matrix of the stylar transmitting tissue in *N. alata* [13]. In addition, some other Trxh can move from cell to cell via plasmodesmata, suggesting an intercellular messenger role as found for some animal Trxs [31–34]. Coincidentally, all the Trxs with a non-cytosolic localisation are clustered within subgroup 2, in which these oxidoreductases are characterised by possessing an N-terminal extension, some also a C-terminal one, whose sequences are not conserved [14].

From the BLASTP results (see above), we identified four Trxh-S2 from *Nicotiana* (S1 Table), namely: NaTrxh (from *N. alata*), NtTrxh2L (*N. tabacum*), NsTrxh2 (*N. sylvestris*) and NtoTrxh2 (*N. tomentosiformis*). All four of them possess identical Nβ sequences (Fig 4).

To include more Trxh-S2 sequences in the analysis, a second BLASTP search was performed using the whole NaTrxh sequence, from which the AtTrxh2 sequence was retrieved (Fig 4), which corresponds to the mitochondrial Trx*h*-2 from *A. thaliana*. This sequence was used for an additional BLASTP. Only Trxh-S2 were considered for a multiple alignment analysis using the *E. coli* Trx1 (EcTrx1) as an outgroup (Fig 4). We also included the mitochondrial PtTrx*h*2, as well as AtTrxh9 and AtTrxh8 because they are known to be membrane-associated proteins [35].

As shown in Fig 4, all sequences contained the conserved redox-active site WCGPC, some contain a C-terminal extension –the longest one belonging to NaTrxh–, and all contain an evident N-terminal extension of similar length when compared to the *E. coli* Trx. Most of the sequences retrieved from the BLASTP search contain similar Nβ sequences but also present differences. All those containing Ser-5 and Ser-9 (positions refer to the original Nβ sequence from Fig 1A) are clustered together (Fig 4, yellow cluster) and, according to our data, might be predicted as secretory proteins. The AiTrxh2 (a Trxh-S2 from *Arachis ipaensis*) was not included in this same cluster because it contains Ala-9. Due to this factor, it can be predicted that AiTrxh2 has a cytoplasmic localisation in concordance with the location of Nβ(S9A)-GFP (Fig 3C). The light-green cluster of Fig 4 contains Trxh-S2 proteins with shorter Nβ sequences. This group exhibits significant residue variations, namely: S5A and S7A, E10D and P11D (positions refer to the original Nβ sequence). This suggests a cellular localisation other than the extracellular space.

The Nβ(S8E)-GFP variant is accumulated in the cytosol when expressed in onion epidermal cells (Fig 3C). A group of sequences was found to contain Glu at this position and, except for one, they have Ala-5 instead of Ser-5; these all were grouped with the mitochondrial localised PtTrx*h*2 (Fig 4, purple cluster), raising the possibility that these differences result in formation of a transit peptide to direct the proteins to the mitochondria. Our data suggest that this short sequence might be an evolutionary source for different cellular localisations. Different combinations of amino acid residues in the N-terminal extension could have evolved to target Trxs to different cellular compartments and organelles, including the apoplast, where they would reduce different target proteins, which amplify the role of these oxidoreductases in several physiological plant processes. This, in turn, would provide an additional explanation of the great diversity of these proteins in plants, particularly among type *h* Trxs.

### The Nβ motif of secretory proteins is predominantly located towards the N-terminus and a structural trait appears to make it function as a secretion signal

The hallmark feature of Trxh-S2 proteins is that they possess N-terminal extensions whose sequences are quite variable [14]. The Nβ motif is located between positions 17 and 27 in NaTrxh (Fig 1A), and when it is fused to the GFP N-terminus (Nβ-GFP fusion protein), it leads to GFP secretion from plant cells [15]. These data suggest that the Nβ motif must be located towards the N-terminal to exert an SP-like role, similar to a hydrophobic typical SP. Therefore, we classified the output sequences considered in Fig 1B –and S1 Table– according to the region in which the Nβ motif is located within each protein. For this aim, each sequence was divided into four equal parts (P1: first 25% of the primary structure towards the N-terminus; P2 and P3: second and third quarter, respectively; P4: the final quarter towards the C-terminus) and was classified according to which quarter the Nβ motif is located in. As shown in Fig 5A, 101 proteins contained the Nβ motif in P1, 65 in P2, 45 in P3 and 88 in P4, resulting in a non-random distribution (χ^2^=27.974, df=3, *p*<0.0001), indicating that indeed, most of the proteins contain this motif towards the N-terminus.

**Fig 5.**
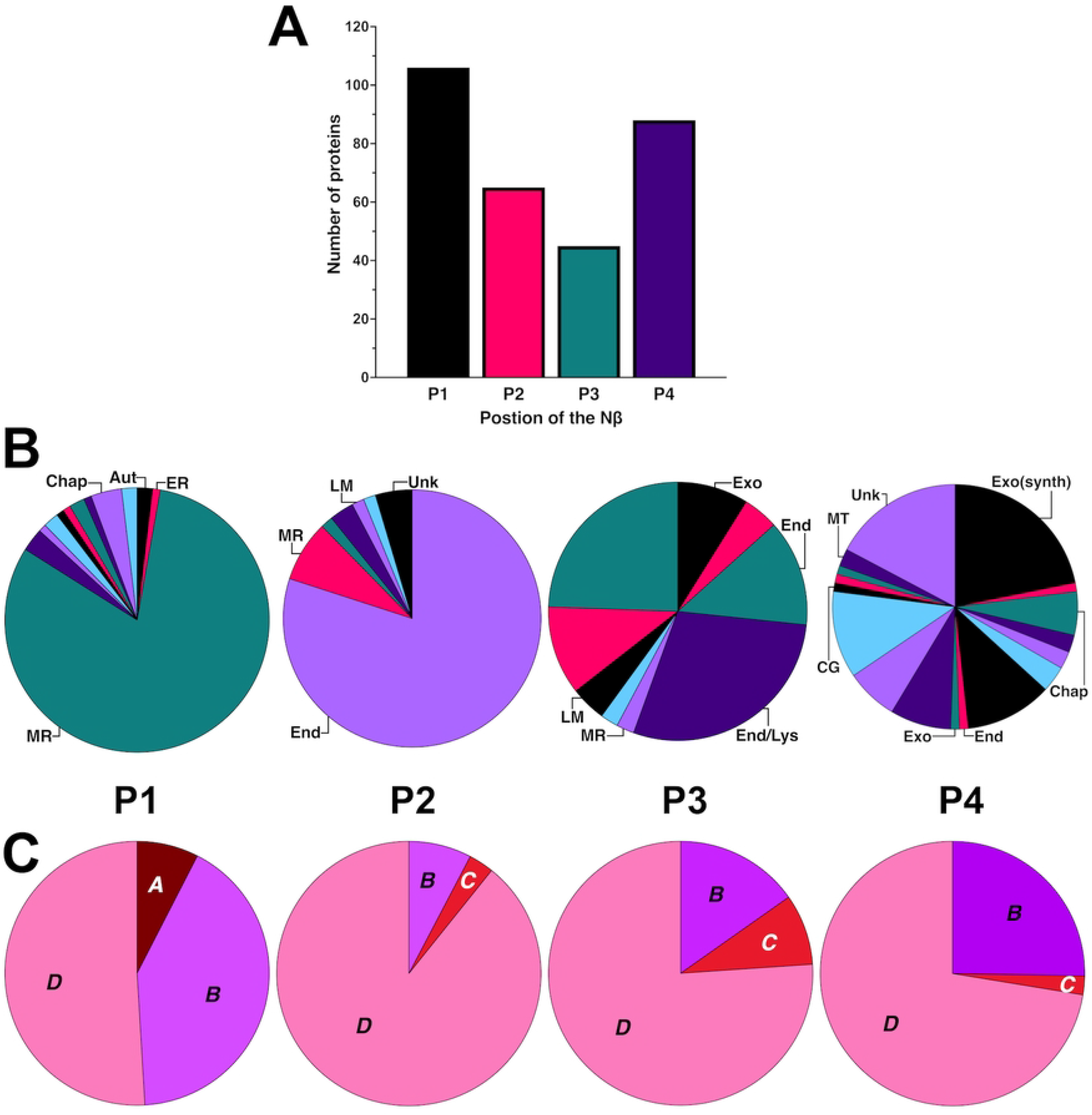
Nβ motif is mostly located towards the N-terminus of extracellular proteins. (A) Distribution of the proteins according to the localisation of the Nβ motif. The sequences were divided into four quarters: P1: 25% of the protein towards the N-terminus; P2 and P3: second and third quarters, respectively; P4: 25% of the protein towards the C-terminus. (B) Annotated functions of the proteins that contain the Nβ motif in each position. Chap: chaperones; Aut: autophagy; ER: endoplasmic reticulum; MR: membrane recycling; LM: lipid metabolism; End: endocytosis; Lys: lysosome; Exo: exocytosis; MT: membrane transport; CG: cell growth; Exo(synth): exocytosis (synthesis); Unk: unkown. (C) Distribution of the cellular localisation based on the categories generated using the *UniProtKB* database in each position (colour code is the same as in Fig 4A).

When each position category was related to the function of each protein, as shown in Fig 5B, P1 and P2 were more homogeneous regarding their annotated roles [81.13% are involved in membrane recycling (P1) and 80% in endocytosis (P2)], in contrast with P3 and P4, which lacked any predominant reported role.

While all the *A* category proteins shown in Fig 3A contain the Nβ motif in P1, none from the category *C* proteins did. Instead, for category *C* proteins, the Nβ sequences were distributed in P2, P3 and P4 (Fig 5C). These data together led us to hypothesise that the Nβ must be located towards the N-terminus of the protein to function as an SP-like sequence.

To test this hypothesis, we generated two different constructs (Fig 6A): (1) the chimeric protein NaTrxhΔNαβ-Nβ-GFP, formed from the core of NaTrxh without its N-terminal extension (NaTrxhΔNαβ) followed by the Nβ-GFP fusion sequence to assess a central position of the Nβ motif; and (2) GFP-Nβ, which served to evaluate the role of the Nβ motif when located in the C-terminal position.

**Fig 6.**
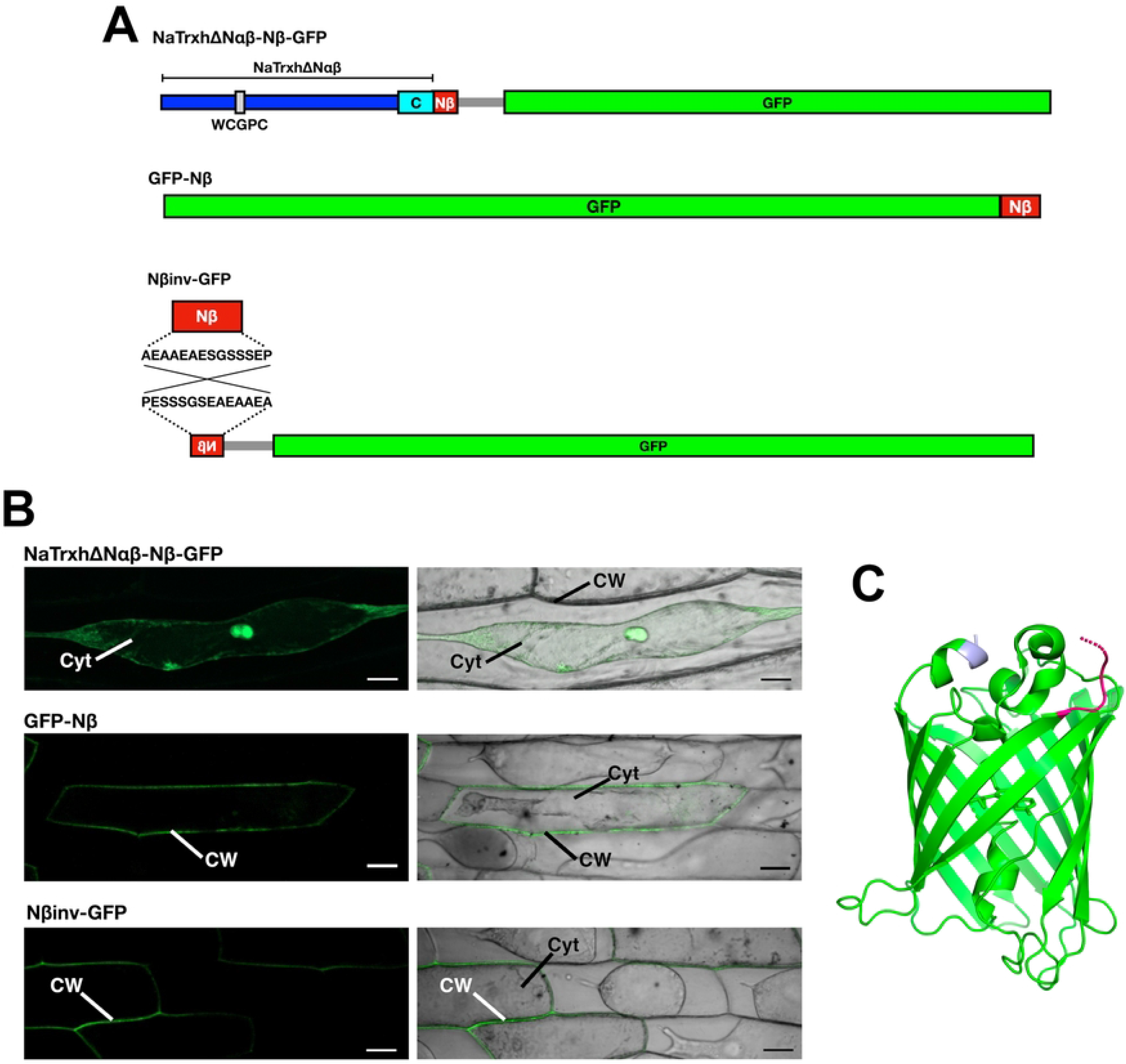
Nβ motif recognition is probably due to a structural feature rather than a positional one. (A) Schematic representation of the constructs generated to assess the position of the Nβ motif as a secretion signal and with the Nβ sequence inverted towards the GFP N-terminus. (B) Transient expression assays in onion epidermal cells of the NaTrxh(ΔNαβ)-Nβ-GFP chimera, the GFP-Nβ and the Nβ(inv)-GFP fusion proteins as in Fig 2. Cyt: cytoplasm; CW: cell wall. Scale bar: 50 μm. (C) Tertiary structure of GFP (PDB 6YLQ) indicating its N-(pink) and C-termini (gray). The figure was generated using PyMOL.

When the NaTrxhΔNαβ-Nβ-GFP chimeric protein was transiently expressed in onion epidermal cells, GFP fluorescence, as predicted, was observed in the cytoplasm (Fig 6B), indicating that the Nβ motif was not identified as a secretion signal within the middle core of the protein, which corresponds to P2 and P3 positions of our previous analysis (Fig 5). Unexpectedly, when the Nβ motif was localised at the C-terminus (GFP-Nβ construct), the GFP signal was detected in the apoplast of the onion epidermal cells (Fig 6B), indicating that the Nβ motif is able to direct secretion whether localised at the N- or the C-terminus. It is noteworthy that, although none of the proteins that contain the Nβ motif towards its C-terminus (P4) are annotated as secretory proteins (*A* category; Fig 5C), in none of the cases is it present at the very C-terminus; the closest Nβ sequence to the C-terminus end of the cytoplasmic proteins (*C* category) is located 122 residues far from it (NP_694967.3 accession code, which is 648 amino acids long; S1 Table).

In the case of the GFP-Nβ protein, the Nβ motif is not only at the very C-terminal end of the protein, but it is also likely to be a mobile element that would be free and exposed to the solvent in a similar manner to the N-terminal extension of NaTrxh [36]. As long as the Nβ motif is free and solvent-exposed, no matter if it localises at the N- or C-terminus, factors –yet to identify– interact with it, directing translocation of GFP into the ER. This latter assumption is based on the tertiary structure of GFP (PDB 6YLQ), in which both N- and C-termini are outside the β-barrel and are oriented towards the same direction (Fig 6C). From the BLASTP analysis, XP_002018569.1 –510 amino acids long– and PLN81894.1 –562 amino acids long– (both GenBank accession codes) contain the Nβ motif at P4 (12 and 15 residues from its C-terminal, respectively) were included in the *B* category (S1 Table). According to the models predicted by AlphaFold in *UniProtKB*, the Nβ motif might be mobile and solvent exposed in both cases, suggesting that these proteins might be secreted. As expected, PLN81894.1 is associated with PM and transport activity (S1 Table). These data reinforce the hypothesis that the *B* category is possibly overrated.

To assess the relevance of the Nβ motif itself, its charge and/or the free and solvent-exposed structure it needs to function as a signal peptide, an inverted sequence version of the Nβ was fused to the GFP N-terminus (Nβinv-GFP; Fig 6A). When the Nβinv-GFP fusion protein was transiently expressed in onion epidermal cells it localised to the extracellular space (Fig 6B) as the Nβ-GFP, GFP-Nβ and NaTrxh-GFP do. This outcome indicates that the Nβ motif’s overall charge –negative at physiological pH– is essential to lead protein secretion rather than the amino acid position. The negative charge is mainly provided by Glu-4 and Glu-10, both present in *A* and *C* category Nβ sequences (Fig 3B). Still, position 9 must contain a serine residue, as shown in Fig 3C, or at least a neutral polar residue to maintain the hydrophilic profile of the Nβ sequence.

### The Nβ-directed secretion involves a post-translational translocation to the endoplasmic reticulum

NaTrxh utilises the ER-Golgi elements of the secretion pathway, as directed by the Nβ motif and demonstrated by the Nβ-GFP fusion protein, which also passes through the ER and the Golgi apparatus [15]. This might suggest a co-translational translocation of the protein towards the ER for its secretion following the conventional pathway. However, the Nβ motif is not hydrophobic, which is a hallmark feature of an orthodox SP, and is required for recognition by SRP during translation, translocating the mRNA-ribosome-nascent protein complex to the ER membrane to further continue translation towards the ER lumen [6,8]. Nevertheless, we discard this scenario for Nβ-directed secretion because Nβ is predominantly hydrophilic.

The fact that the GFP-Nβ protein was found in the extracellular space (Fig 6B) discards the possibility of GFP-Nβ translation coupled to the ER translocation by SRP. Therefore, we hypothesised that the GFP-Nβ protein is fully translated in the cytosol and then translocated to the ER to continue via the Golgi apparatus to ultimately reach the apoplast [15]. To test this, the ER retention signal KDEL was added to the GFP-Nβ [GFP-Nβ(KDEL); Fig 7A] and transiently expressed in onion epidermal cells. As shown in Fig 6B, the GFP-Nβ(KDEL) protein was detected within the cell exhibiting a typical ER-retained protein pattern, similar to that found for NaTrxh-GFP(KDEL) and Nβ-GFP(KDEL) fusion proteins using the same type of expression assays [15]. Thus, our data provide evidence that once the Nβ motif is free and exposed to the solvent in any protein, either at its N- or very C-terminal end, and is at least 8-residue long, is recognised as a transit peptide to the ER to be followed by protein secretion (Fig 8). It is possible that the functional sequence of this signal is X-X-X-E-S/G-G-S-<u>S/E</u>-S-E-P, where the fifth position must be a neutral amino acid residue (at least a serine or a glycine) and position 9 a serine residue. The eighth position needs a serine in the case of plant proteins –especially for Trxh-S2– for a secretory function, but if there is Glu-8 –together with Ala-5– it is possible that the sequence instead functions as a mitochondrial transit peptide, as might be the case for AtTrxh2 [31]. Moreover, in the case of animal systems, there is another possibility: that of a requirement for a glutamate (Glu-8) in order to provide a secretory function. Both possibilities are worthy of further investigation.

**Fig 7.**
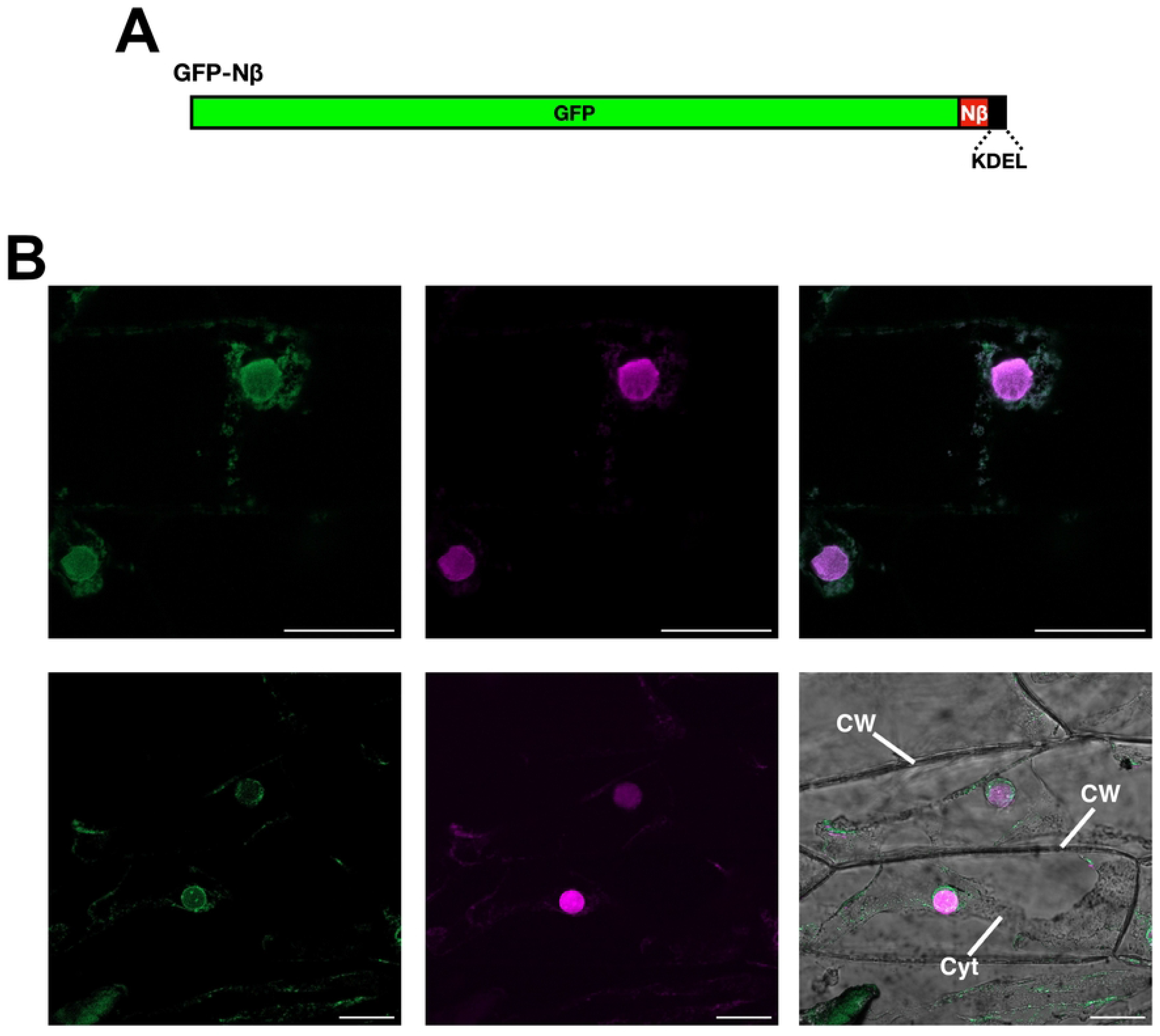
The GFP-Nβ fusion protein uses the ER for its secretion. (A) Schematic representation of the GFP-Nβ construct with the ER retention signal KDEL [GFP-Nβ(KDEL)]. (B) Transient expression assay in onion epidermal cells of the GFP-Nβ(KDEL) protein. Nuclei were stained with propidium iodide before observation (magenta fluorescence). Left panels: GFP fluorescence; middle panels: propidium iodide fluorescence; right panels: GFP and nucleus-labelled merged image. Cyt: cytoplasm; CW: cell wall; Nu: nucleus. Scale bar: 50 μm.

**Fig 8.**
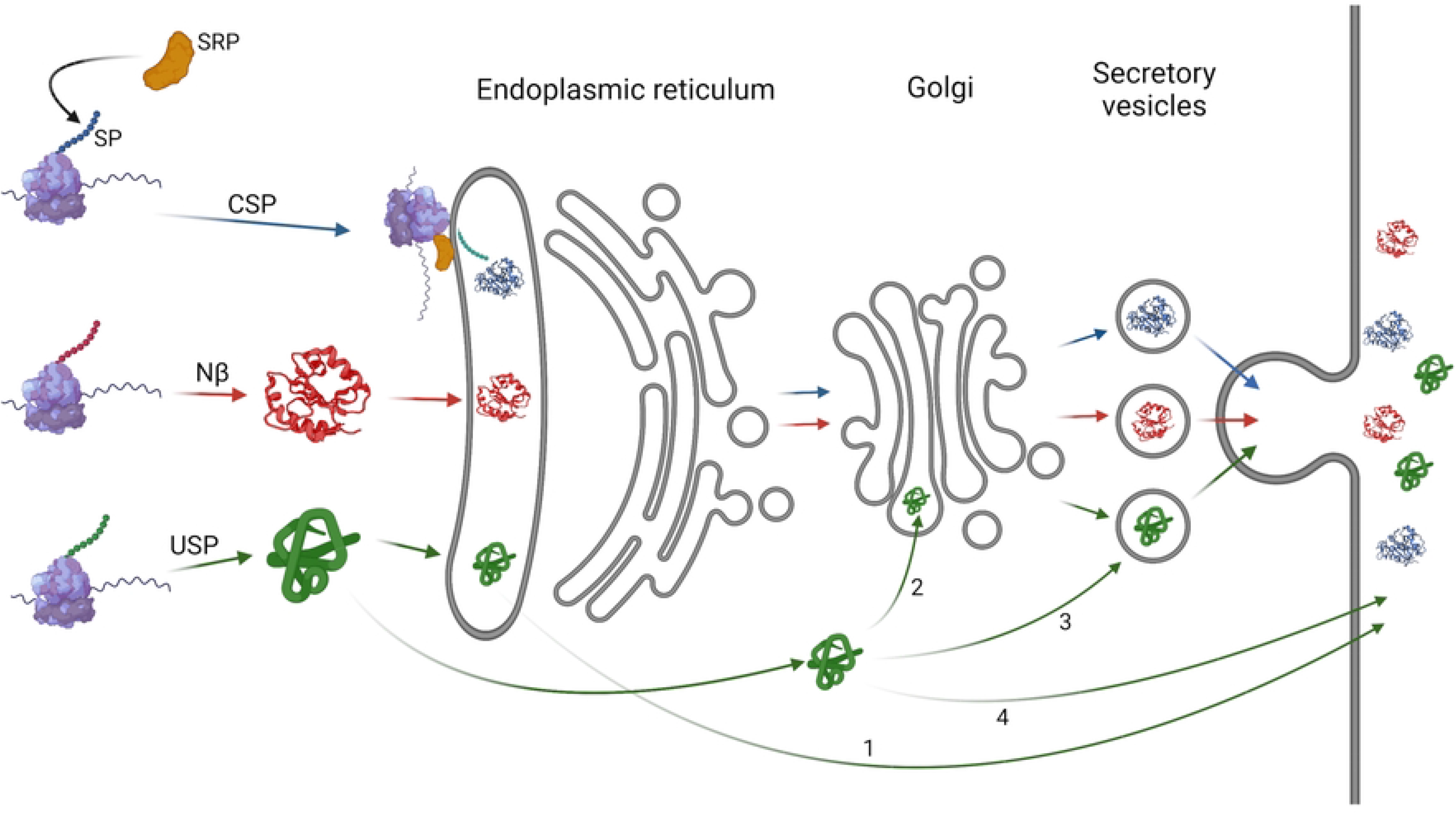
The Nβ “semi-conventional” secretion pathway differs from the conventional and unconventional pathways. In the conventional secretion pathway (CSP), during translation of a protein with a signal peptide (SP), the SP is recognized by SRP, resulting in the translocation of the mRNA-ribosome-nascent protein to the endoplasmic reticulum lumen, where the SP is cleaved, and translation is completed. The unconventional secretion pathways (USP) comprise: Golgi apparatus bypass (1), where the protein is directly secreted from the ER; ER-independent routes, where proteins may follow secretion after Golgi incorporation (2), through incorporation in multivesicular bodies (3), or through alternative direct secretion pathways (4). In the case of Nβ motif-containing proteins, such as NaTrxh, these are translated in the cytosol and, in an SRP-independent manner, are translocated to the ER to then progress to secretion via the CSP elements (ER, Golgi and secretory vesicles). Figure created using Biorender.

## Discussion

The conventional secretion route, in addition to passage through the ER, the Golgi apparatus, and the exit of the proteins from it within vesicles, includes the assistance of co-translational incorporation of the protein –while translated in the ribosome– into the ER by SRP (Fig 8). This translocation is the result of the SRP-SP interaction, in which the SP at the N-terminal end of the protein must be a hydrophobic element [8]. The Nβ motif is quite the opposite since it is hydrophilic –negatively charged– as previously described [15] and further analysed in this work. This eliminates the possibility of Nβ being recognised by SRP. However, the NaTrxh-GFP, Nβ-GFP ([15] and Fig 2B) and GFP-Nβ fusion proteins reach the extracellular space by passing through the ER (Figs 6B and 7B). This, and particularly the latter, in which the Nβ is at GFP C-terminus, indicates that the protein was fully translated in the cytosol and then incorporated into the ER (Fig 8), indicating that the ER localisation was probably not due to an SRP-dependent translocation of the nascent protein during its translation, as occurs in the conventional secretion pathway. The Nβ-led secretion, as reported for NaTrxh-GFP and Nβ-GFP, also involves the Golgi apparatus since it is brefeldin-A sensitive [15]. Therefore, the secretory pathway for Nβ-containing proteins differs from both conventional and unconventional secretion pathways. While it is not ER nor Golgi independent –varying from the unconventional proposed pathways–, the entry to the ER-Golgi route also differs from the conventional one, suggesting an additional pathway that would be “semi-conventional” in nature (Fig 8). Likewise, the secretion of GFP-Nβ protein suggests that the Nβ motif is not processed as occurs with most classical SPs [8]. The presence of this motif in mature NaTrxh is crucial for interaction with the S-RNase in order to regulate its ribonuclease activity [16].

The results reported here suggest that the Nβ motif of NaTrxh acts as a transit peptide towards the ER (Figs 6B and 7B). Some proteins localised to different cell compartments are first translated into the cytosol and then translocated towards the target organelle due to the presence of a transit peptide. However, some remain in the cytosol until a signal directs them to the corresponding organelle. A good example of this latter group are nuclear proteins, such as transcription factors, that remain in a latent form in the cytosol and only in response to certain stimuli are they re-located to the nucleus. An example of this is the NF-κB complex, which contains a nuclear transit peptide but remains in the cytosol because its inhibitor, I-κβ, is tightly bound and obscures the transit peptide [37,38]. During an immune or inflammatory response, for instance, I-κB is phosphorylated and degraded, releasing NF-κB and exposing the nuclear localisation sequence, resulting in its translocation towards the nucleus, where it will activate transcription of specific target genes [37,38]. A similar scenario could occur with NaTrxh or any Nβ-containing protein with the features described here, but translocated towards the ER rather than nucleus. This would represent an efficient mechanism to regulate a portion of soluble protein secretion, which would occur when the Nβ is exposed.

NaTrxh is directly involved in *Nicotiana*’s gametophytic SI system, known as S-RNase-dependent SI [16]. SI is a genetically controlled system that acts as a barrier to self-pollination and involves complex pollen-pistil interactions in which secreted stylar proteins play key roles [17,19,39,40]. The S-RNase –the female S determinant– and NaTrxh colocalise to the extracellular matrix of the stylar transmitting tissue in *N. alata* [13]. S-RNases and other proteins involved in this gametophytic SI system contain a conventional SP and are secreted by the stylar cells [41,42], the exception being NaTrxh, where secretion is due to the presence of its Nβ motif [15].

In either compatible (non-self-pollen) or incompatible (self-pollen) crosses, S-RNase enters the pollen tube cytoplasm [43,44]. For self-pollen rejection, reduction of a disulphide bond in S-RNase by NaTrxh is required in order to increase its ribonuclease activity; transgenic *Nicotiana* plants expressing a non-functional NaTrxh are unable to reject wild-type self-pollen [16]. These data indicate that secreted stylar NaTrxh must enter the pollen tube cytoplasm, where the electron donor system is available, in order to reduce S-RNase.

Once in the pollen tube cytoplasm, how does NaTrxh, with its Nβ motif, avoid secretion from the pollen tube? How does it stay in the cytosol and not become targeted to the ER? The answer could rely on the stable interaction between S-RNase and NaTrxh, in which both N- and C-termini are involved [13,15,16]. Although this interaction has only been observed *in vitro*, the N-terminal extension of NaTrxh is essential to reduce S-RNase [16]. The NaTrxh-S-RNase interaction results in a protein-protein complex where the Nβ motif is stabilised, as modelled using the crystal structures of both NaTrxh (PDB 6X0B) and S_F11_-RNase (PDB 1IOO), where stabilising interactions are detected between the NaTrxh N-terminal extension and S_F11_-RNase [36].

If not completely hidden, the NaTrxh N-terminal extension might become a stable and ordered element [36]. This evidence also explains why proteins, including the NaTrxhΔNαβ-Nβ-GFP chimera, that contain the Nβ in an inner position, although potentially exposed to the solvent, are not secreted, because this motif might form stable secondary structural elements. Additionally, in the case of the GFP-Nβ protein, the GFP C-terminus is likely to be accessible, away from the GFP β-barrel. The Nβ motif would be mobile, disordered and fully exposed to the solvent, which might be the reason it works as an ER translocation and further secretion signal in the GFP-Nβ fusion protein (Figs 6 and 7). Trxs are widely distributed proteins, from prokaryotes to eukaryotes, that reduce disulphide bonds of target proteins [45]. The polymorphism of these proteins reflects the different roles they are involved in, which are as wide as the target proteins they reduce [22,46]. Plant Trxs represent a large and complex system in which 8 classes of them have been identified. The type *h* thioredoxins form the largest and most heterogenous group and are subdivided into three subgroups; the subgroup 2 Trxh contains extensions towards the N-terminus some also at the C-terminus [14,29]. Although the Trxh-S2 N- and C-terminal extensions are not fully conserved, some share common features. Those Trxh-S2 proteins that have the most similar sequence to the Nβ of NaTrxh were clustered together (Fig 4), and possibly they all are secreted in a similar manner. However, another group of Trxh-S2 contained some differences. They grouped with a known mitochondrial Trx, AtTrx*h*2 [31], as shown in Fig 4. This reinforces the idea that this short sequence functions as a transit peptide, but interestingly, select changes to the amino acid sequence –at least two are detected in this work– might influence its target cellular localisation. Therefore, having initially acquired an Nβ sequence, further evolutionary changes to the sequence could lead to different Trx homologs being differentially localised in the cell, where they could reduce new target proteins. For example, whilst having serine residues at positions 5 and 8 results in a secretory motif, Ala-5 and Glu-8 might result in a mitochondrial localisation sequence.

Dissecting both NaTrxh extensions has provided useful information on how they are involved in NaTrxh cellular localisation and its specificity towards its identified target protein in *N. alata* styles –the S-RNase–. Both *E. coli* Trx and NaTrxh reduce insulin disulphide bonds as expected from any Trx [13,46], but *E. coli* Trx is not able to reduce S-RNase [16]; the major difference between these two Trxs is the occurrence of the N- and C-terminal sequences in NaTrxh. In addition, according to the structural model of the NaTrxh-S-RNase protein complex, the NaTrxh N-terminus assists the correct orientation of the interaction to reduce only one of the four disulphide bonds that the S-RNases typically contain [36]. The differences detected in the Nβ sequences of the Trxh-S2 (Fig 4), apart from those in positions 5, 8 and 9, might contribute to different specificities regarding their respective target proteins. However, there is also the Nα motif, which in the case of NaTrxh, contributes to the interaction with S-RNase and its reduction [15,16]. This motif, with some variation, is also present in the other analysed Trxh-S2 (Fig 4) and could contribute to their target specificity too.

Finally, finding other leader-less proteins containing a similar Nβ motif towards the N-terminus –like NaTrxh– with annotated functions related to protein traffic, membrane mobility and secretion (some experimentally confirmed) raises the possibility that their cellular localisation is due to this short sequence. Therefore, the Nβ motif sequence might help to predict the protein localisation of those proteins that contain it.

The cellular localisation of some proteins containing a similar Nβ motif is not yet known (*B* and *D* categories of our analysis; Figs 3 and 5). Some, particularly the Trxh-S2 proteins, might move from the *B* or *D* category to the *A* category, as was the case of NaTrxh itself. Notably, the Nβ motif of NaTrxh and other plant proteins containing this motif, must contain a serine at position 8. However, a glutamate residue was usually found at this position in animal-secreted proteins (Fig 3). This opens the possibility of distinctive functional secretion motifs between plant and animal proteins that are worthy of being investigated.

## Materials and methods

### *In silico* analysis, image processing and statistics

A BLASTP query with default settings (NCBI; http://www.ncbi.nlm.nih.gov) was executed using the Nβ motif sequence as a query. The output sequences were retrieved and classified according to the type of organism they belonged to (animals, fungi, plants). Sequence logos were constructed using the WebLogo server (http://weblogo.threeplusone.com/) to show the consensus sequences [48,49].

Confocal images were processed with ImageJ Fiji. Statistical analysis and graphs were developed using GraphPad Prism.

### Deletion of 3 and 6 positions on the Nβ motif and generation of GFP fusion DNA constructs

The *NaTrxh* sequence was previously cloned into pENTR/D-TOPO (Invitrogen) [15] to generate pENTR:NaTrxh. This was used as a polymerase chain reaction (PCR) template to amplify and produce [Platinum SuperFi PCR master mix (Invitrogen)] the *NaTrxhΔNα(+3)* and the *NaTrxhΔNα(+6)* sequences using the forward primers Nβ(-3)-F or Nβ(-6)-F, respectively (Table S2), with the NaT-R reverse primer for both (S2 Table).

Using the pEG103:Nβ construct [15], in which the *Nβ* sequence is upstream of the *GFP* gene in the pEarleyGateway103 vector [50] as template, the *Nβ(-3)-GFP* and the *Nβ(-6)-GFP* sequences were obtained by PCR with the same forward primers as above and the GFP-R as reverse (S2 Table). Additionally, we also amplified the full *Nβ-GFP* sequence from the pEG103:Nβ construct using the Nβ-F and GFP-R primers (S2 Table).

PCR products were cloned into pENTR/D-TOPO (Invitrogen) following the manufacturer’s instructions. Then the sequences were transferred by recombination using LR recombinase enzymatic mixture (Invitrogen) to the pEG103 vector [*NaTrxhΔNα(+3)* and *NaTrxhΔNα(+6)*, generating the pEG103:NaTrxhΔNα(+3) and the pEG103:NaTrxhΔNα(+6) constructs, respectively], or to pK2GW7 [51] in the case of *Nβ-GFP*, *Nβ(-3)-GFP* and *Nβ(-6)-GFP*, resulting in the pK2:Nβ-GFP, pK2:Nβ(-3)-GFP and the pK2:Nβ(-6)-GFP constructs, respectively. All constructs were transformed into *Escherichia coli* XL10-GOLD cells (Agilent), pEG103 construct transformants were selected using kanamycin (50 μg/mL in solid and 100 μg/mL in liquid medium) and pK2GW7 construct transformants with spectinomycin (50 μg/mL). All constructs were confirmed by DNA sequencing.

### Site-directed mutagenesis

To obtain the different Nβ variants (S5G, S8E, S8D, S8A and S9A; positions corresponding to the Nβ motif sequence), the pK2:Nβ-GFP construct was used as template for site-directed mutagenesis by PCR using the QuickChange Lightning Site-Directed Mutagenesis Kit (Agilent) following the manufacturer’s instructions with the concomitant mutagenic primers for each target base substitution (S2 Table). Presence of the desired mutations was confirmed for each construct by DNA sequencing and confirmed no other nucleotide changes were detected. All resulting sequences were transformed into *E. coli* XL10-GOLD cells and selected using spectinomycin (50 μg/mL) resistance.

### Generation of DNA constructs with the Nβ motif at different positions, with the Nβ sequence inverted, or with the ER retention signal

To assess the relevance of the Nβ position within the protein to its role as a secretion signal sequence, we generated two different constructs: (1) NaTrxh(ΔNαβ)-Nβ-GFP; and (2) GFP-Nβ. The first one, encodes a chimeric protein that contains the Nβ motif between the core of NaTrxh (without its N-terminal extension) and GFP, and the second one contains this motif at the GFP C-terminus.

The *NaTrxh(ΔNαβ)-Nβ* sequence was generated by two sequential PCR amplifications. In the first one, the pair of primers NaTΔαβ-F and NaT-Nβ-R1 (S2 Table) were used with the pENTR:NaTrxh construct [15] as a template. The PCR product was analysed by electrophoresis and purified to use it as the template for the second amplification, in which the same forward primer was used and the reverse one was NaT-Nβ-R2 (S2 Table).

The *GFP-Nβ* sequence was also generated by two sequential PCR amplifications. For the first one, with the pEG103 vector as the template, the primers GFP-F and GFP-Nβ-R1 (S2 Table) were used. The PCR product was analysed by electrophoresis and purified to use it as template for the second amplification, in which the same forward primer was used and the reverse one was GFP-Nβ-R2 (S2 Table).

The *Nβinv-GFP* sequence was obtained after two sequential PCR amplifications as described for the *NaTrxh(ΔNαβ)-Nβ* and *GFP-Nβ* sequences but using pK2:Nβ-GFP as the template and the primers Nβinv-F1 and GFP-R2 (S2 Table) in the first reaction and for the second amplification the primers Nβinv-F2 and GFP-R2 were used (S2 Table).

The GFP-Nβ(KDEL) protein contains the ER retention signal at its C-terminus. Its coding sequence was generated by PCR using the pENTR4:GFP-Nβ construct (described below) as the template and the primers GFP-F and NβKDEL-R (S2 Table).

The four sequences were subcloned into pJET using the CloneJET PCR cloning Kit (ThermoScientific) and confirmed by DNA sequencing. The inserts were released by *Bam*-HI and *Eco*-RI digestion and ligated to pENTR4 (Invitrogen; previously digested with the same restriction enzymes), separated by electrophoresis and purified (eliminating the dual selection cassette between the *att*L1 and *att*L2 sites). The resulting constructs [pENTR4:NaTrxh(ΔNαβ)-Nβ, pENTR4:GFP-Nβ, pENTR4:Nβinv-GFP and pENTR4:GFP-Nβ(KDEL)] were transformed into *E. coli* XL10-GOLD cells, selecting transformant cells with kanamycin (50 μg/mL in solid and 100 μg/mL in liquid medium).

To generate the *NaTrxh(ΔNαβ)-Nβ-GFP* fusion, we transferred the sequence from pENTR4:NaTrxh(ΔNαβ)-Nβ to pEG103 by LR recombination. Since both entry and destination vectors provide kanamycin resistance, we followed a previously reported strategy [52] of amplifying by PCR [using pENTR4:NaTrxh(ΔNαβ)-Nβ as the template] a fragment from upstream of the *att*L1 site (pENTR4-F primer; S2 Table) to downstream of the *att*L2 site (pENTR4-R primer; Table S2), purifying the resulting PCR product and using it for the recombination step. The pENTR4:GFP-Nβ, pENTR4:Nβinv-GFP and pENTR4:GFP-Nβ(KDEL) were used directly for the LR recombination reaction into pK2GW7, which confers spectinomycin resistance (selected using 50 μg/mL spectinomycin). All these constructs [pEG103:NaTrxh(ΔNαβ)-Nβ, pK2:GFP-Nβ, pK2:Nβinv-GFP and pK2:GFP-Nβ(KDEL)] were transformed into *E. coli* XL10-GOLD cells.

### Transient expression assays in onion epidermal cells

Each construct, either in pEG103 or pK2GW7, was individually bombarded into onion epidermal cells as previously reported [15] using 2 cm^2^ slices, which were placed on 0.5x Murashige-Skoog medium, pH 5.5-57, with 1 % (w/v) agar and 4.5 g/L TC-gel (Caisson). After 24-48 h after particle bombardment, onion epidermal cells were incubated in 1.0 M NaCl for 5 to 10 min to separate the plasma membrane from the cell wall (plasmolysis). Fluorescence was visualized using an Olympus FV 1000 confocal microscope or Zeiss LSM 800 confocal microscope [From Fig 3C: Nβ(S5G), Nβ(S8A), Nβ(S8D) and Nβ(S9A)]; in both cases with 485/670 nm excitation/emission light for GFP. Where indicated, the nuclei were stained with propidium iodine (Sigma-Aldrich) prior to plasmolysis and visualised with 570/670 nm excitation/emission light.

## Acknowledgements

We thank K. Jiménez-Durán (USAII-FQ-UNAM), R. Rincón-Heredia and A. Rosas-Arellano (IFC-UNAM) for their technical assistance with confocal microscopy. To M. Olvera-Flores (FQ-UNAM) for her technical assistance with bombardment transformation. To A. Beacham (Harper Adams University) for the editing work. We also thank Posgrado en Ciencias Biológicas, UNAM, for the support provided to A.Z-G. and J.A.J-D.

---

**Table S1.** Sequences raised from the BLASTP analysis. Retrieved sequences from the BLASTP analysis with an identical or similar Nβ motif sequence. Protein sequences are arranged in rows with the information of each one of them in the columns. Column A: assigned number of the proteins, organised according to the taxon each one corresponds to. Column B: GenBank accession code; Column C: name of the protein and the species where it is from; Columns D and E: covery and identity percentages with Nβ motif, respectively; Column F: annotated functions; Column G and H: species and taxon where each protein is from, respectively; Column I: amino acid position of the Nβ motif in the primary structure; Column J: position where the Nβ motif locates within the primary structure (P1, P2, P3 or P4, according to the analysis described in the main text); Column K: size of the primary structure; Column L: cellular localisation. EC: extracellular, Cyt: cytoplasmic, NA: data not available; Column M: score regarding the cellular localisation according to the *UniProtKB* database, where 5 is the highest and 1 the lowest; Column N: score resulted from SignalP 6.0 to predict the presence of a signal peptide, where 1 is a high probability of the presence of a signal peptide; Column O: organelle or cellular region to which the protein is associated (NA: data not available). The colour code represents the different categories generated in this analysis (described in the main text): green (*A* category), yellow (*B* category), red (*C* category) and gray (*D* category).

**Table S2.**
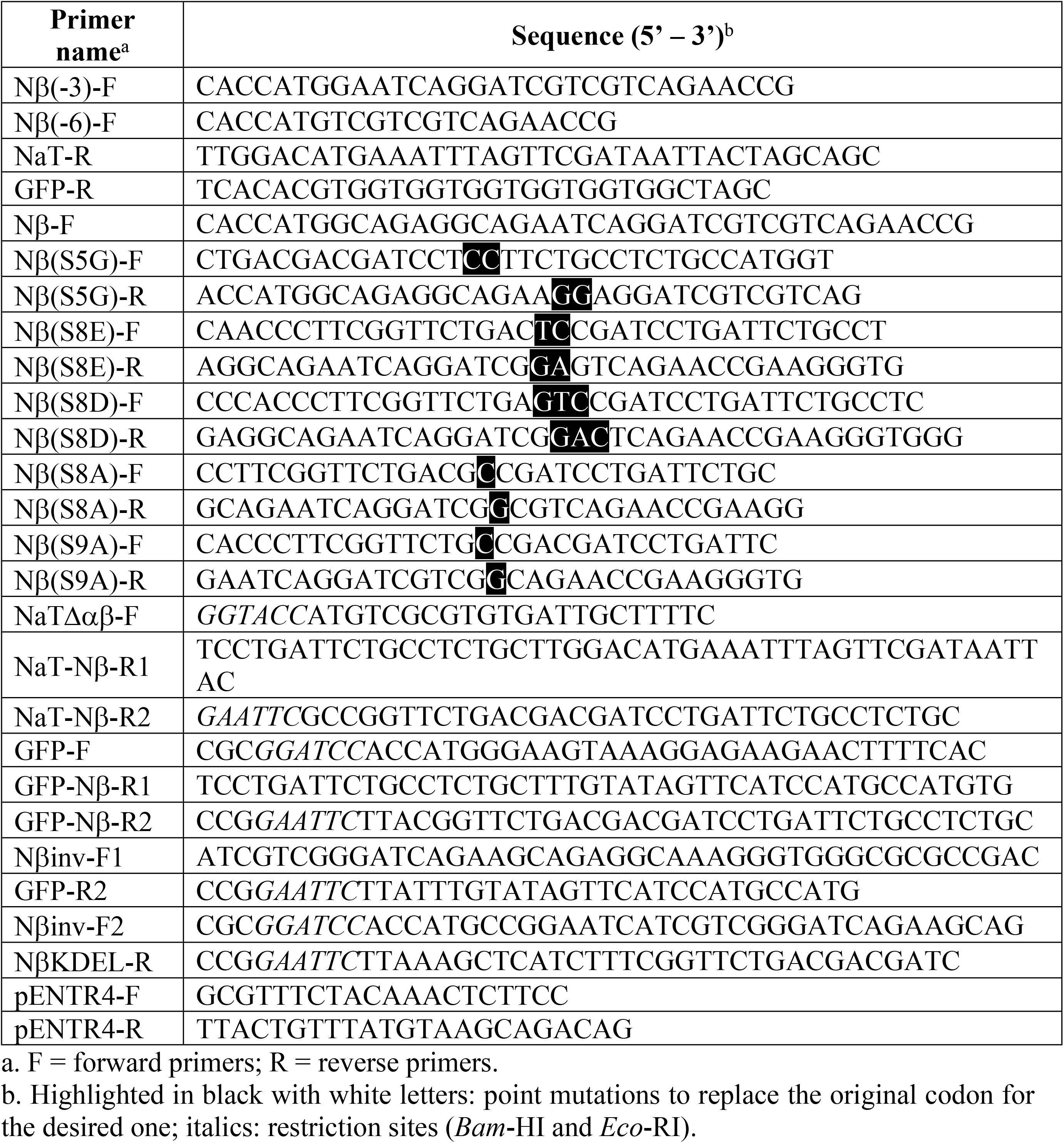
Primers used for the different constructs generated for transient expression assays in onion epidermal cells.

